# Raspberry plant stress detection using hyperspectral imaging

**DOI:** 10.1101/2023.02.22.529512

**Authors:** Dominic Williams, Alison Karley, Avril Britten, Susan McCallum, Julie Graham

**Affiliations:** James Hutton Institute, Invergowrie, Dundee; James Hutton Limited, Invergowrie, Dundee

## Abstract

Monitoring plant responses to stress is an ongoing challenge for crop breeders, growers and agronomists. The measurement of below ground stress is particularly challenging as plants do not always show visible signs of stress in the above ground organs, particularly at early stages. Hyperspectral imaging is a technique that could be used to overcome this challenge if associations between plant spectral data and specific stresses can be determined. In this study, three genotypes of red raspberry plants grown under controlled conditions in a glasshouse were subjected to below ground biotic stresses (root pathogen *Phytophthora rubi* and root herbivore *Otiorhynchus sulcatus*) or abiotic stress (soil water availability) and regularly imaged using hyperspectral cameras over this period. Significant differences were observed in plant biophysical traits (canopy height and leaf dry mass) and canopy reflectance spectrum between the three genotypes and the imposed stress treatments. The ratio of reflectance at 469nm and 523nm showed a significant genotype-by-treatment interaction driven by differential genotypic responses to the *Phytophthora rubi* treatment. This indicates that spectral imaging can be used to identify variable plant stress responses in raspberry plants.

## Introduction

One of the major challenges facing agriculture is the task of increasing food production while also reducing inputs applied to crops. Pests and diseases reduce crop yields and quality of the harvested product, and timely, effective identification and control is therefore needed. Integrated pest management (IPM) approaches are becoming increasingly important in reducing reliance on chemical pesticides^[1]^. Use of resistant crop varieties is the basis for IPM, along with crop surveillance and precision monitoring so that controls can be applied at an early stage of pest or disease establishment ^[2]^. High throughput monitoring of crops is, therefore, an important component of IPM, both for pest and disease monitoring and for large-scale crop phenotyping in breeding programmes to detect genotypes with resistance or tolerance. Plant imaging, and in particular hyperspectral imaging, provides a method for non-invasive monitoring of large plant numbers^[3]^ to detect crop responses to pests and pathogens.

The feasibility of using hyperspectral imaging to detect diseases has been tested in several crops ^[4,5]^. Hyperspectral imaging for plants is normally carried out in the visible (400-700nm), near infra-red (700-1,000nm) and short wave infra-red (1,000-2,500nm) regions of the spectrum. Hyperspectral images contain information for hundreds of wavelength bands covering several regions of potential interest. It differs from standard red-green-blue (RGB) images by collecting far more detailed colour information from the wavelength bands, and from point-based spectroscopy by also containing spatial information^[6,7]^. The methods of image capture vary including using cameras attached to unmanned aerial vehicle (UAV) platforms which fly over crops^[8]^, motorised ground-based platforms passing through or over the crop canopy^[6,7]^, handheld measuring techniques including both imaging and non-imaging spectrometers^[9]^, and imaging of collected plant samples under controlled lighting conditions^[10]^.

Raspberry is a high value fruit crop with increasing global consumption. In raspberry cultivation, major root pathogens/pests affecting plant health include the root rot pathogen *Phytophthora rubi* and root-feeding larvae of the vine weevil *Otiorhynchus sulcatus*, both of which require prompt detection of infection or infestation to prevent serious crop losses. While effort has been invested in breeding resistant varieties, the detection of suitable genetic markers for breeding could be accelerated by using high throughput methods for accurate phenotyping in field conditions. Our previous work using a raspberry mapping population has shown that hyperspectral imaging can be used to detect genotypic differences in plant physical characteristics ^[11]^. This study demonstrated that certain hyperspectral imaging wavelengths and ratios were heritable and were associated with genetic markers on linkage groups (QTLs), including some that co-located with QTLs for physical crop characteristics. Similarly, hyperspectral imaging of grapevine leaves has been shown to be capable of distinguishing crop varieties indicating that hyperspectral imaging can provide an alternative method for phenotyping fruit crops^[12,13]^.

The development of hyperspectral imaging for rapid non-invasive assessment of pest and pathogen damage in raspberry has great potential to benefit the soft fruit industry. Hyperspectral tools have previously been tested for disease detection in tomato crops. One study used a combination of UAV and lab-based imagery to detect disease from an early stage of pathogen infection^[12]^, although two diseases with similar visual symptoms were not readily distinguished. Similar work in citrus fruit^[13]^ for detection of canker found that signs of disease could be detected using either UAV imaging or benchtop imaging of samples. In grapevine, static hyperspectral imaging was used to detect infection by grapevine vein-clearing virus ^[14]^. Machine learning techniques were then used to build a classifier capable of distinguishing infected and uninfected plants.

To date, these studies represent the few examples where hyperspectral imaging has been applied for stress detection in perennial crops. Raspberry provides a suitable model for testing the utility of spectral signatures, building on our existing knowledge of spectral signals for genotype differentiation^[11]^, to identify genotypic responses to biotic and abiotic stress. In this study, experimental manipulation of plant stress was carried out using three contrasting plant genotypes under controlled (glasshouse) conditions to test plant responses to reduced water availability, root infection with the pathogen *P. rubi* and root infestation with herbivorous vine weevil larvae. Data was collected on plant growth, physiology, health, and final biomass as well as plant imaging data collected every two weeks. Our aim was to identify spectral signatures of plant stress in raspberry by assessing the relationship with biophysical measures of plant performance. This was achieved with the following three objectives: i) analysing imaging data to identify wavelengths and wavelength ratios that respond differentially to the stress treatments and show genotype-by-stress treatment interactions; ii) analysing biophysical data to assess plant performance under control and stress treatments; and iii) testing the relations between these spectral signals and biophysical measures of plant performance to identify any wavelengths or ratios that co-vary as potential early indicators of plant stress.

## Methods

### Experimental material and stress treatments

96 raspberry plants were grown in pots with a standard compost mix for raspberry. Three varieties were used, Latham, Glen Moy and 0946/12. Latham is a North American variety that is known to be resistant to *Phytophthora rubi*, Glen Moy is a variety developed at the James Hutton Institute that is susceptible to *Phytophthora rubi*, and 0946/12 is an advanced selection from the James Hutton Limited breeding program containing a *Phytophthora rubi* resistance genetic marker^[15]^. Four treatments were used: low water availability, high water availability, a disease treatment (root rot, *Phytophthora rubi*) and a pest treatment (vine weevil, *Otiorhynchus sulcatus*). All plants were watered using an automated irrigation system which watered the different treatments independently. Soil moisture probes were used to control the irrigation system with irrigation being triggered when soil moisture level fell below a particular value: for the low water treatment, irrigation was turned on at 18% volumetric soil moisture content (m^3^/m^3^); for all other treatments, the irrigation was turned on at 32% soil moisture (m^3^/m^3^). This moisture level was derived from taking the mean of six pots in the treatment excluding the lowest and highest reading to prevent the system being skewed by extreme readings. The *Phytophthora rub*i treatment was applied by placing *Phytophthora rubi* (4 plugs of culture grown on nutrient agar plates) in the pots adjacent to the plant roots. The vine weevil treatment was applied by placing 12 eggs of vine weevil in the pots in a small indent of the substrate adjacent to the plant roots; eggs were collected from insect cultures maintained on excised strawberry leaves at 20°C with 16 h daylength. The plants were arranged in 4 blocks each comprising two rows of 12 plants (24 plants in total) using a split plot design, with treatment assigned at random to plots within each block, so that plants receiving the same irrigation level were grouped together. Within each plot, two replicate plants of each genotype were assigned at random to positions, giving a total of eight replicate plants per genotype-by-treatment combination. A diagram of the set up can be seen in supplementary materials.

### Biophysical Plant Traits

A series of biophysical plant and soil characteristics were monitored weekly. The number of leaves on each plant were counted. The height of the canopy of each plant from the base of the pots was measured in cm. Leaf health was recorded with a plant level visual 1-5 score where 1 was a healthy undamaged leaf and 5 was a dead leaf. Towards the end of the experiment a visual 1-5 overall plant health score was also recorded in the same way as leaf health.

Leaf chlorophyll content was estimated using a hand-held Chlorophyll meter (CCM-200: Opti-Sciences, Tyngsboro, Massachusetts, USA), which provides a chlorophyll content index (CCI) for a 0.71 cm^2^ area of leaf based on absorbance measurements at 660 and 940 nm. Meter readings were converted to total chlorophyll concentrations (chlorophylls a and b, in μg per unit area of leaf) using the equations of Lichtenthaler and Wellburn^[17]^ to construct a calibration curve for representative leaf discs extracted in 80% acetone. Measurements were taken from a single leaf of each plant every two weeks.

Stomatal conductance was measured using an AP4 porometer (Delta T devices, Cambridge, United Kingdom). This uses a humidity sensor to measure the volume of water released from the stomata of a leaf. Measurements were taken from a single leaf of each plant every two weeks.

Soil moisture measurements were taken using a handheld soil moisture meter (SM150, Delta T devices, Cambridge, United Kingdom). This was inserted into the soil from the surface measuring the soil moisture content of the surface layer (*c*. 5 cm depth) of the pots. Measurements were taken in each pot every two weeks.

At the end of the experiment, plants were harvested by cutting at the stem base and separating stem and leaf material. Roots were gently washed to remove the compost substrate. Total leaf area was measured in cm^2^. Fresh and dry mass for each plant were recorded for leaves, stem and root. A 1-5 score of root damage was assigned where 1 was a healthy undamaged root and 5 a badly damaged root. For the root rot treatment, root mass was not quantified as root material had to be destroyed to prevent pathogen escape. For the vine weevil treatment, the number and fresh mass of vine weevil larvae in the substrate was recorded when the roots were harvested.

### Hyperspectral Imaging

We used a hyperspectral imaging platform developed at James Hutton Institute ^[16]^. This has a visible near infrared red (VNIR) scanner which covers the wavelengths 400-950nm with 394 spectral bands. The imager is a pushbroom line scanner meaning movement is needed between the camera and object of interest to capture the image; in this case the imaging platform was moved along each block/row of plants. The imager was mounted on a hand pushed trolley which gathered images horizontally at around 0.5m from the plants.

The plants were imaged in 8 rows of 12 plants (one row per image) every week for the final 12 weeks of the experiment. At the start of each row a white reference tile was placed and captured in the image. Exposure was adjusted to ensure that this tile was not overexposed. Once the exposure was set the platform was manually pushed at a slow walking pace along the row. At the end of the row the platform was reversed, and a second set of images taken of the plants. Each image contained a lateral view of 12 plants comprising half of the block. As each block comprised two rows, the scanner was turned through 180-degrees to image the second row relative to the first row. Figure 1 below shows a false colour image generated by the scanner.

**Figure 1:**
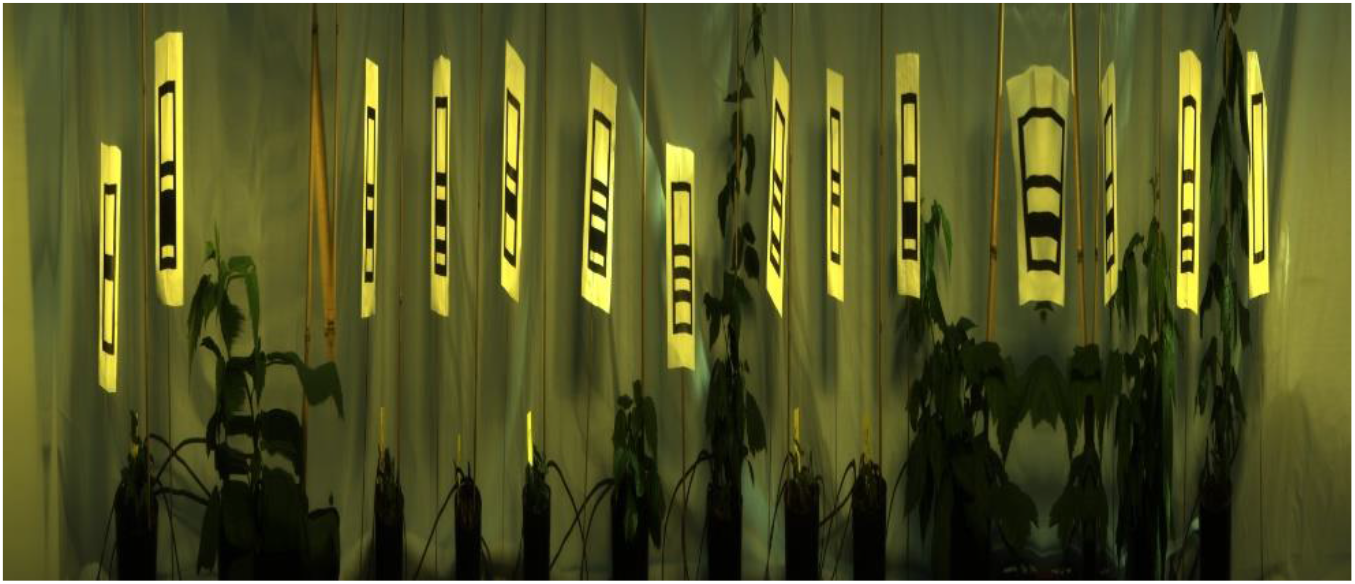
Example image generated by scanners. Showing 12 plants in a lateral view.

### Image and Spectral Data Analysis

To analyse the data from the hyperspectral camera an image analysis pipeline was developed. Plant material was identified by applying a threshold to the normalised difference vegetation index (NDVI). NDVI was defined as NDVI = (IR – red)/(IR + red) where red was defined as the mean intensity between the wavelengths of 650 to 680 nm, and IR as the mean intensity between the wavelengths of 710 to 740 nm. The white reference tile was manually selected in the image. The images were then split into individual plants by identifying gaps between the plants as columns in an image where no plant pixels were detected. A manual inspection and correction step was needed to deal with cases where the plants had grown close together in the glasshouse. Following the segmentation and splitting stages, a mean spectrum for each plant was generated. Tests were carried out only using some of the plant pixels for analysis: pixels were categorised according to their height in the image and by the mean intensity or brightness across all wavelengths. It was found that using the brightest 10-25% of pixels produced the most significant difference between treatments so only these pixels were used to generate the mean value for each plant. The thresholds were set on a per plant basis so the same proportion of pixels from each plant was used. This is most likely due to these pixels representing consistently illuminated surfaces of the plants.

To carry out further analysis of the data, four different types of measures were used. The first was the individual wavelengths, the second was a series of wavelength ratios that had been gathered from the hyperspectral imaging literature (table 1), the third was a series of ratios generated from our previous research with raspberry ^[11]^. Finally, the top ten principal components were calculated using all the data for each date (see ‘Statistical analysis’ below) and were analysed since they provided a simple summary of the highly correlated wavelength variables.

**Table 1:**
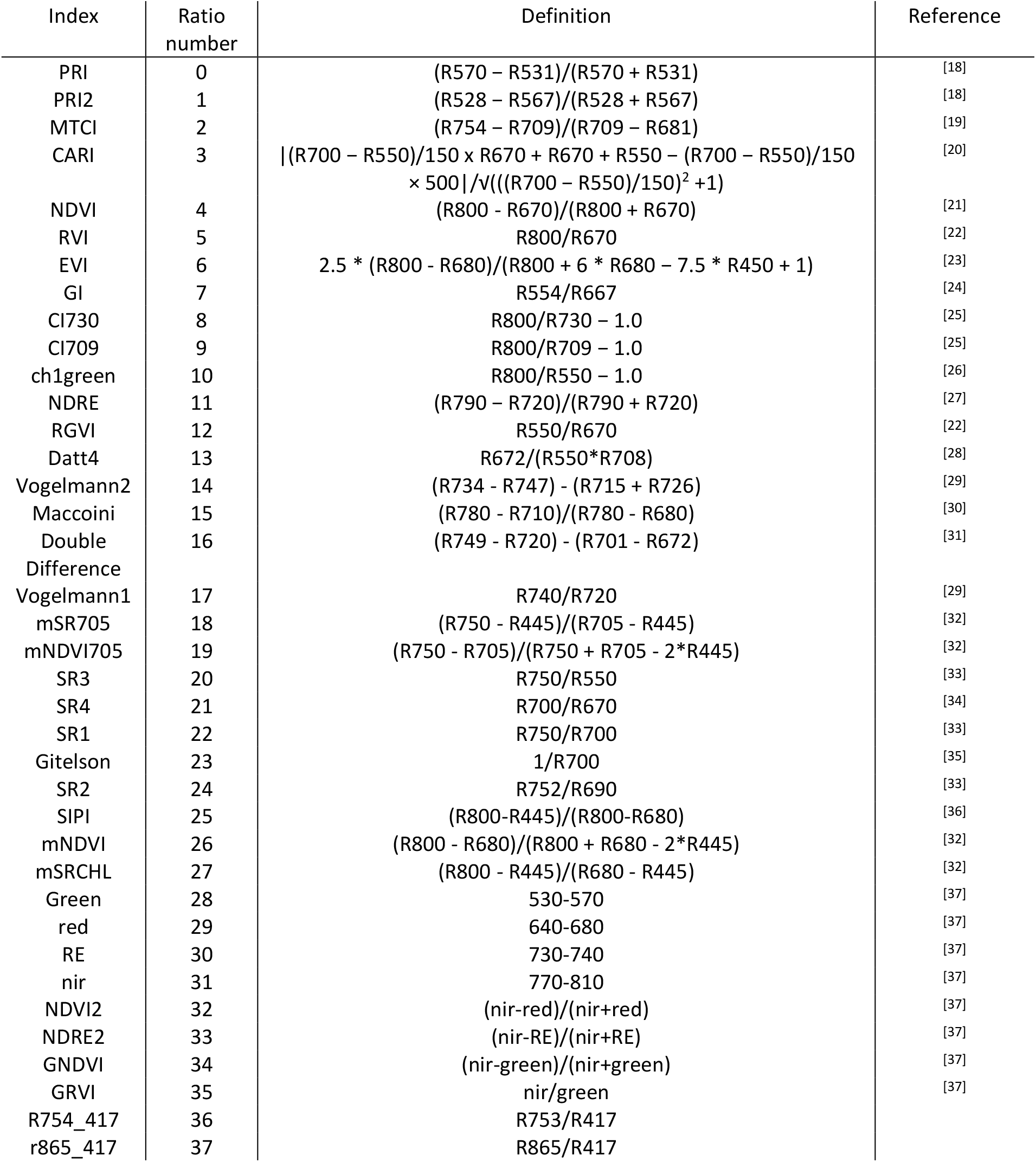
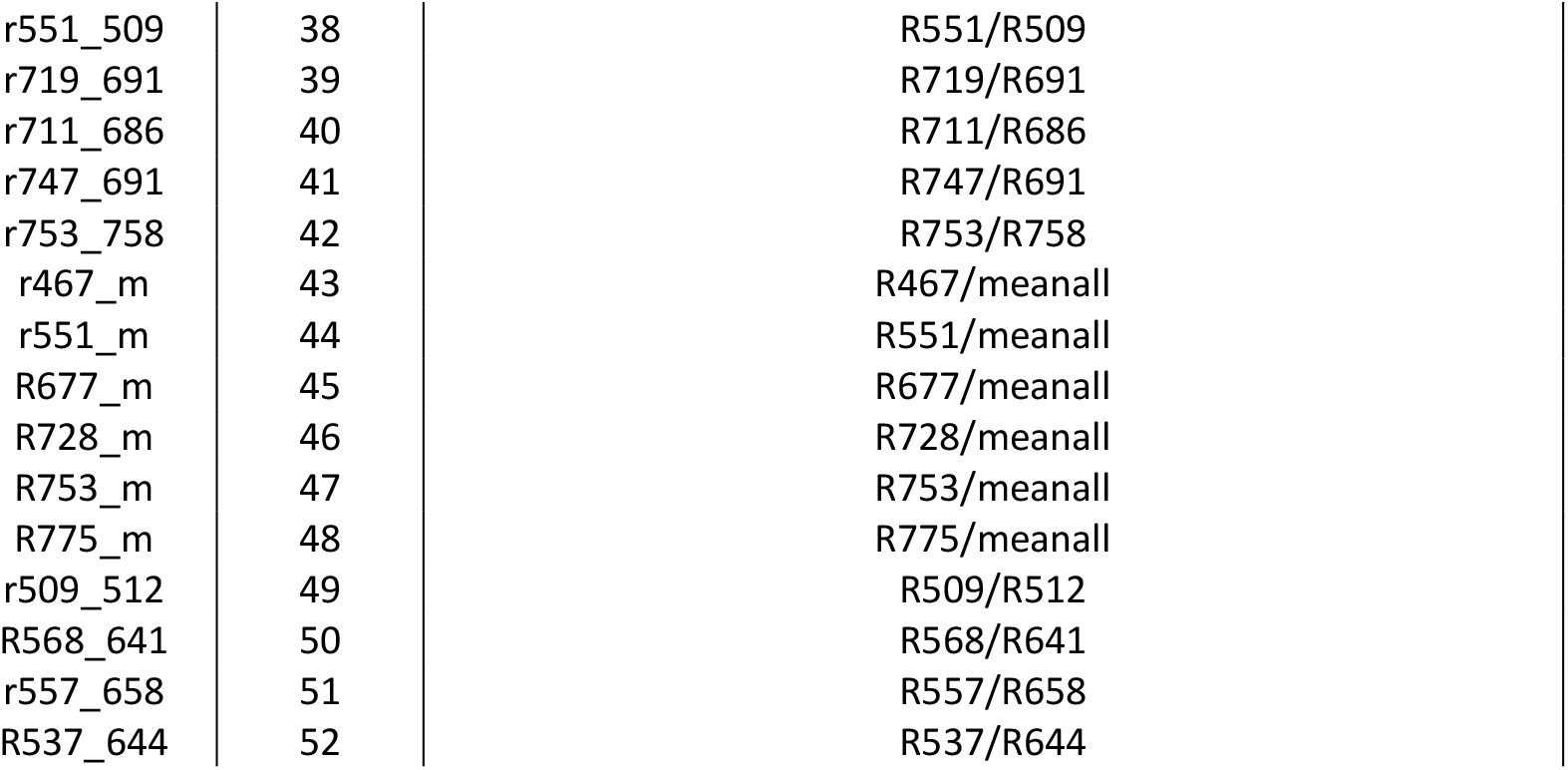
List of all wavelength ratios used in this study. Rxxx indicates band at xxx nm wavelength. meanall is mean intensity of all wavelengths across image. References are included for the ratios taken from the published literature.

### Statistical analysis

The initial glasshouse plan had originally been designed as a split-plot design with four blocks. During the imaging it was established that each block needed two images and there were significant camera position effects between these two images (which were taken at a 180-degree angle to each other). This meant in practice we had an 8-block design rather than the 4 blocks originally planned. The position of each of the different genotypes within each treatment plot was randomised and consequently each genotype/treatment combination was not equally represented in each of the 8 blocks. To counter this problem, a residual maximum likelihood (REML) method was used to determine if there were significant genotype, treatment or genotype*treatment fixed effects, and image number (one to eight) was used as a random effect in the model. Individual wavelengths and a series of selected ratios and principal components of imaging data were examined for significant differences. The principal components were calculated independently for each date using Genstat (20^th^ edition) using the sum of square and products method. Genstat 20^th^ edition was used to carry out the statistical analysis and the analysis was carried out independently for each date.

The biophysical data was analysed taking into account the 4 block split-plot design using ANOVA to determine if there were significant genotype, treatment or genotype*treatment fixed effects, and block number (one to four) was used as a random effect in the model. This was carried out in Python using the statsmodels package^[38]^.

A comparison was made between the spectral data and biophysical measurements using a correlation analysis in Python. Pearson correlation coefficient was calculated between the biophysical data and the spectral data collected on the closest available date. Data from the final imaging date was compared with the post-harvest measures.

To establish that the stress treatments had been imposed successfully, their effects on different biophysical measures was assessed, with these measures ranked in the following order according to whether they reflected long- or short-term stress responses. More weight was given to measures that show long term effects of stress (No. leaves per plant, canopy height, leaf and root dry mass, root damage score, plant health score, leaf chlorophyll content) rather than short term stress indicators (soil moisture, stomatal conductance).

The leaf damage score was used primarily to monitor other sources of stress that can arise in a glasshouse environment such as mildew, red spider mite and thrips infestations.

## Results

### Biophysical Plant Traits

For ease of presentation, the biophysical data are split between the continuous measures that were carried out throughout the experiment and those measured at the end of the experiment. Figure 2 shows the significance of genotype, treatment and genotype-by-treatment interactions in the continuous monitoring data. Several of the measures, including leaf number, canopy height, health score and soil moisture, consistently showed significant treatment effects at a p<0.01 threshold in the later stages of the experiment. Only chlorophyll content did not respond significantly to the treatments. Leaf number, canopy height and plant health scores were smallest in the low water treatment, reflecting the low soil moisture availability. Conversely, leaf porometer readings were highest in the high-water treatment (Supplementary Tables 1). Vine weevil and *P. rubi* treatments tended to be intermediate in their effects. This indicates we were successful in applying stress treatments to the plants. Significant genotype effects were seen for all the different measures on most weeks confirming the genotypes examined exhibited contrasting growth and other physiological characteristics.

**Figure 2:**
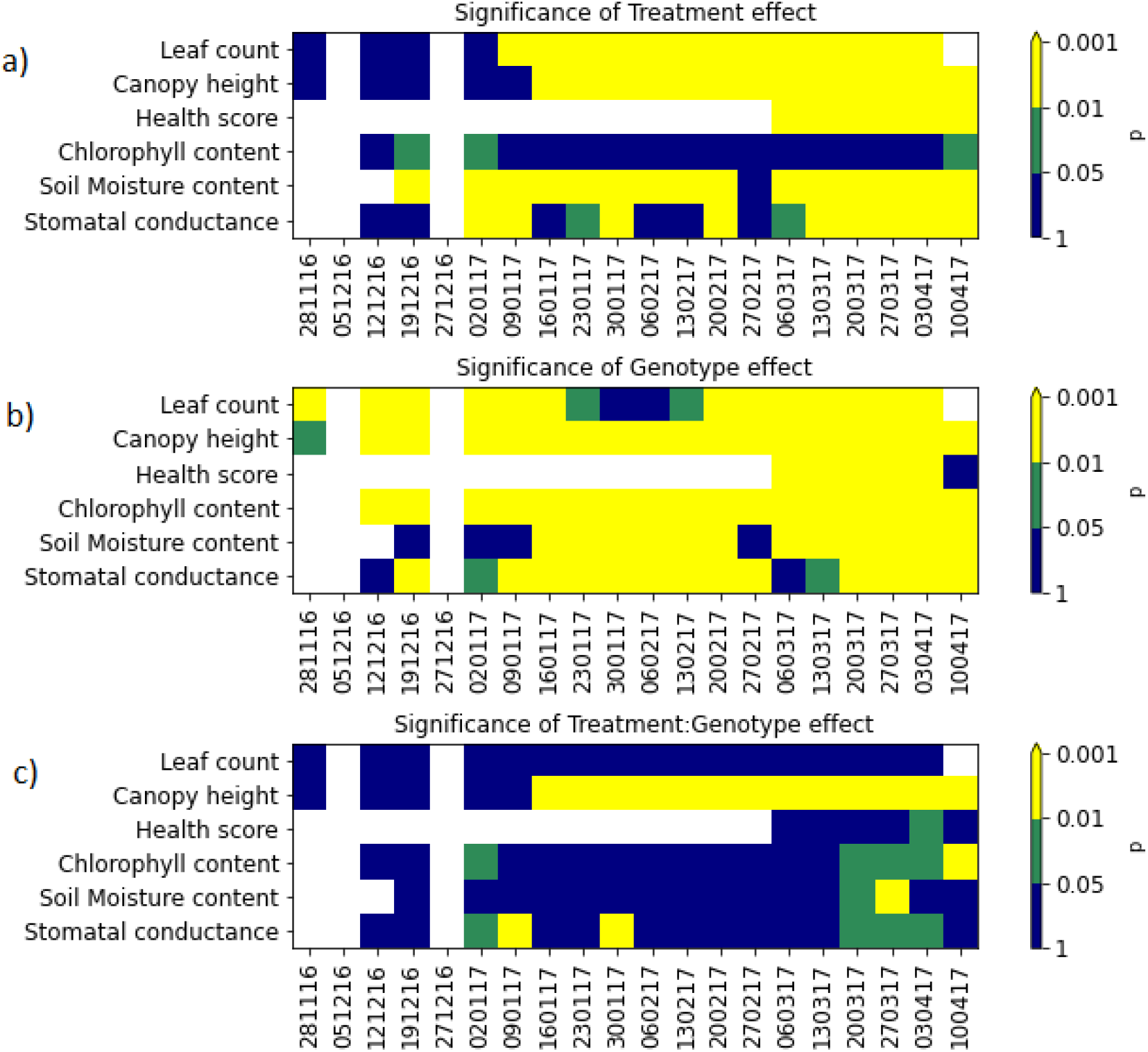
Heat maps showing significance of treatment, genotype and interaction effects for plant growth and other biophysical measurements during the experimental period. White indicates no data, blue not significant at p=0.05 threshold, green indicates significant at p=0.05 threshold but not 0.01 and yellow indicates significant at p=0.01 threshold.

Significant treatment*genotype interactions were seen consistently in canopy height, but fewer significant interactions were observed for other growth traits. For canopy height Glen Moy and 0946/12 were taller than Latham and both were tallest in the *P. rubi* treatment, while Latham canopy height was tallest under the vine weevil treatment; the results from the final week of the experiment are shown in figure 3a below. Leaf chlorophyll content showed significant interaction effects in the final few weeks of the experiment, and significant interaction effects were observed for stomatal conductance on some dates. For leaf chlorophyll content, the genotype*treatment interaction was driven by differential genotype responses to the *P. rubi* treatment with Glen Moy and 0946/12 showing their lowest values of chlorophyll content under this treatment but Latham showing the highest value in this treatment. The significant interaction for stomatal conductance was also driven by the effects of the *P. rubi* treatment with Glen Moy showing higher stomatal conductance under this treatment than other treatments, while the other genotypes did not respond as strongly to the *P. rubi* treatment. This indicates that canopy height is a good measure to look at plant*genotype interactions. Plants may still produce similar numbers of leaves but could be smaller when under stress, which might not be detected in leaf count or chlorophyll content measurements.

**Figure 3:**
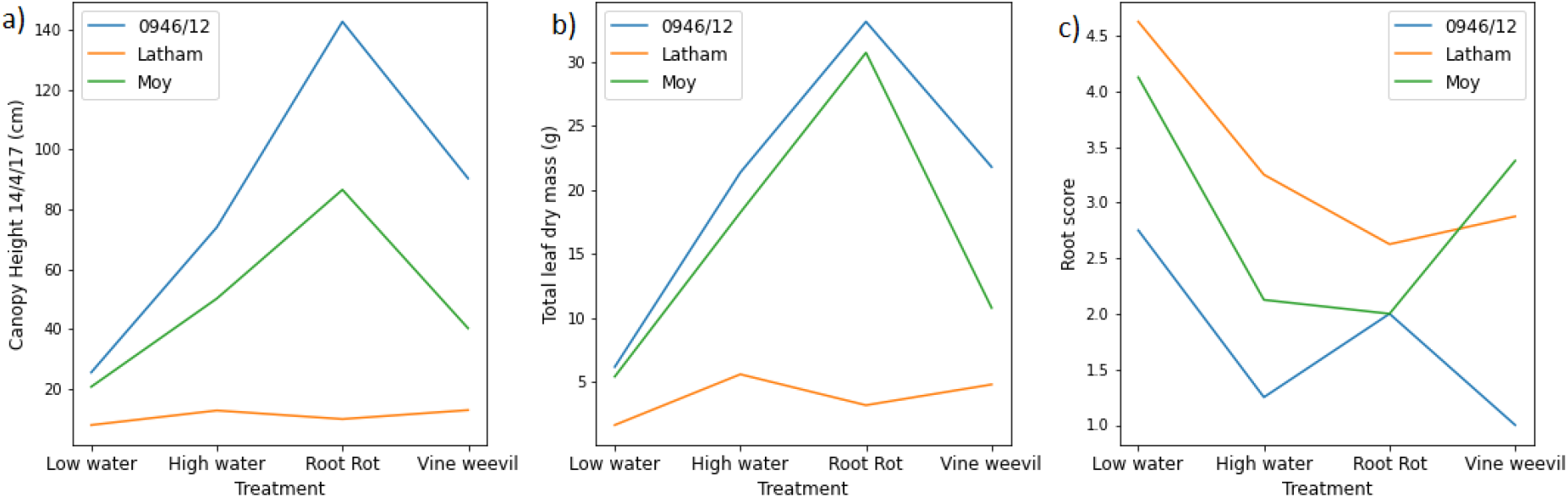
Genotype treatment interaction plots for three biophysical measures of plant performance. There are 8 replicates for each genotype treatment combination. a) Canopy height just before harvest b) total leaf dry mass and c) root damage score

Statistical analysis of plant growth data collected at plant harvest is shown in supplementary figure 1. All measures apart from root length were significant for both treatment and genotype. All measures apart from leaf area and leaf specific mass also showed significant interaction effects. The genotypes 0946/12 and Moy had a larger total leaf mass than Latham, and both had the highest leaf mass in the *P. rubi* treatment. In the vine weevil treatment, Glen Moy developed a lower leaf mass compared with the high-water treatment whereas 0946/12 exhibited similar values in those two treatments. Latham had lower leaf mass than the other genotypes, and the high-water treatment had highest mass of all treatments, as seen in figure 3b. The root damage score results (shown in figure 3c) showed a different pattern. This is a damage score so 1 is a healthy root and 5 is badly damaged. The rank order of the genotypes was similar to canopy height and leaf dry mass, with 0946/12 showing the least damage. In the vine weevil treatment, however, Glen Moy suffered the most damage and 0946/12 the least.

This supports the conclusion that we were successful in applying stress treatments to the plants and that there were differential genotypic responses to stress.

Root damage was scored in the vine weevil and *P. rubi* treatments and were only analysed for genotype responses. There was no significant difference between genotypes in the *P. rubi* damage score which was unexpected due to the contrasting cultivar root rot susceptibilities. For vine weevil, there was a significant difference between genotypes for vine weevil damage score but not for the number or mass of larvae, indicating differences in the plant response to herbivory but not in the level of vine weevil infestation. 0946/12 had the lowest vine weevil damage score of 1 indicating very little damage and Glen Moy had the highest score with a mean of 3.4. Root damage scores showed a similar but inverse pattern to root mass when comparing the treatments indicating it is a good proxy trait for root mass.

We observed unpredicted effects of the *Phytophthora rubi* treatment on the three genotypes. In this treatment, Glen Moy (very susceptible variety) and 0946/12 (the root rot resistance marker genotype) produced largest leaf dry mass, while low scores for root damage were observed for Latham (resistant) and Glen Moy (susceptible). Mean values of all measures can be seen in supplementary table 1.

### Hyperspectral plant traits

The mean reflectance values for each genotype and treatment for a representative date are shown in figure 4.

**Figure 4:**
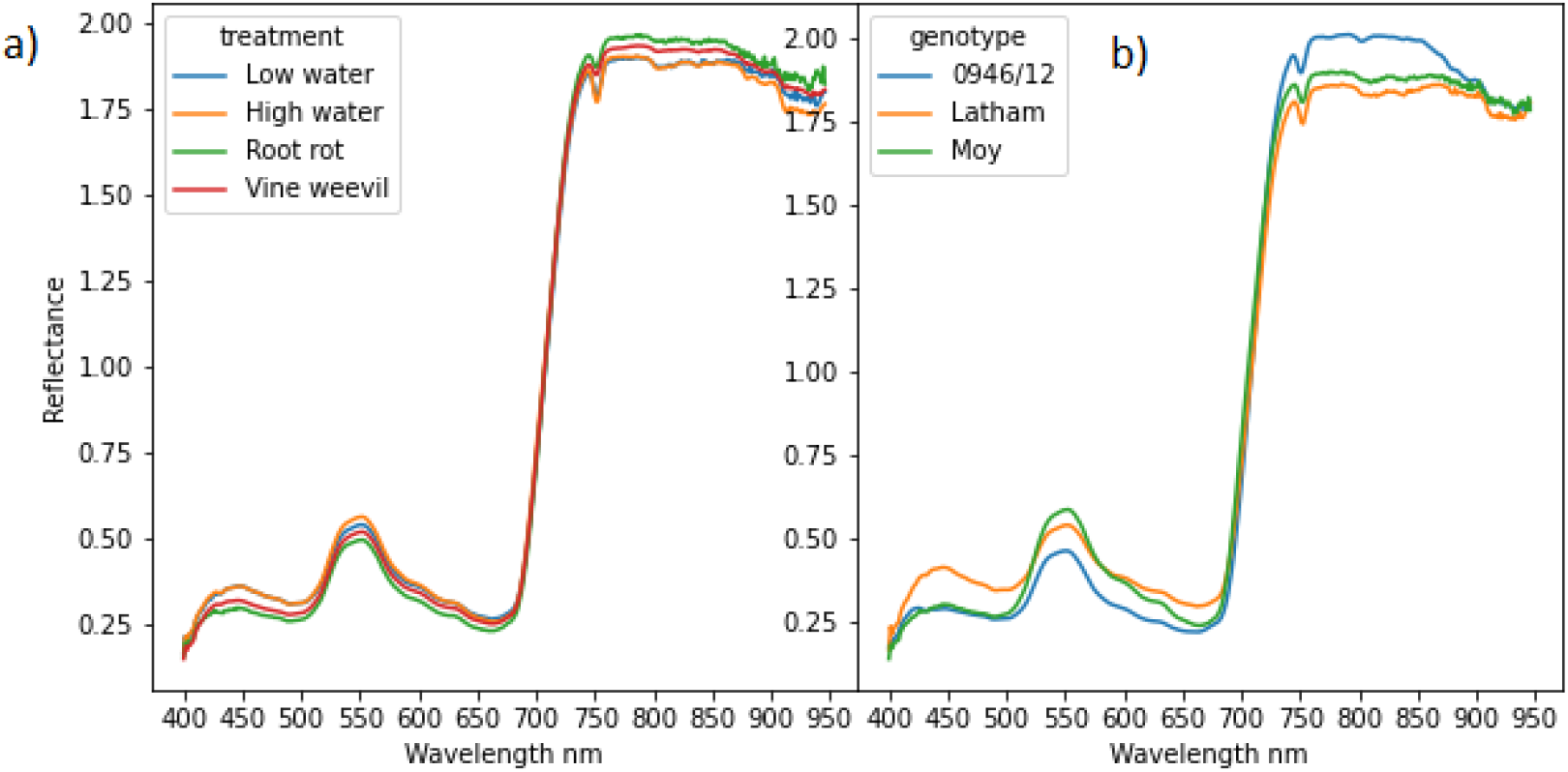
Plot showing mean reflectance of all plants grouped on a) left by treatment and on b) right by genotype. Data were collected on 1/3/17. Wavelengths in the range 400-650nm are in the visible light region, and the peak around 550nm is in the green region. Above 700nm is a steep rise in reflectance which is characteristic of photosynthetically active plant material which absorbs light mainly in the visible region.

The reflectance spectrum of the raspberry leaves followed a similar pattern in the different treatments and genotypes. All plots show a small peak in a region around 550nm which is green light and a sharp increase at around 690nm in the near infra-red. This is the characteristic reflectance profile of photosynthetically active plant material. Larger differences were seen between the genotypes in both visible and near infra-red regions. 0946/12 showed higher reflectance in the infrared while Latham showed higher reflectance in the visible regions. The differences between treatments were smaller than between genotypes, with increased reflectance in the infra-red region in the *Phytophthora rubi* (root rot) treatment and highest reflectance in visible regions for the high water and, to a lesser extent, the low water treatments.

Figure 5 indicates the significance of treatment effects on a series of selected ratios and principal components. The significance effects of genotype on spectral reflectance are not presented here as these have been established in our recent study of field-grown plants ^[11]^. Numerous significant treatment-by-genotype interactions were seen. The most consistently significant interaction was for r469_523 (Ratio 52 on figure 5). This shows a significant treatment-by-genotype interaction for every sampling date from 17/2/17 to the end of the experiment apart from two dates (1/3/17 and 9/3/17). This interaction was driven by differences between the *P. rubi* treatment and the other three treatments, as seen in figure 6. Towards the end of the experiment more ratios showed a significant interaction effect, which is consistent with an expected increase in the level of stress experienced by plants in the later stages of the experimental treatments, indicating that these wavelengths could be used as indicators of plant responses to stress. Several ratios also showed significant treatment effects, particularly on 23/2/17.

**Figure 5:**
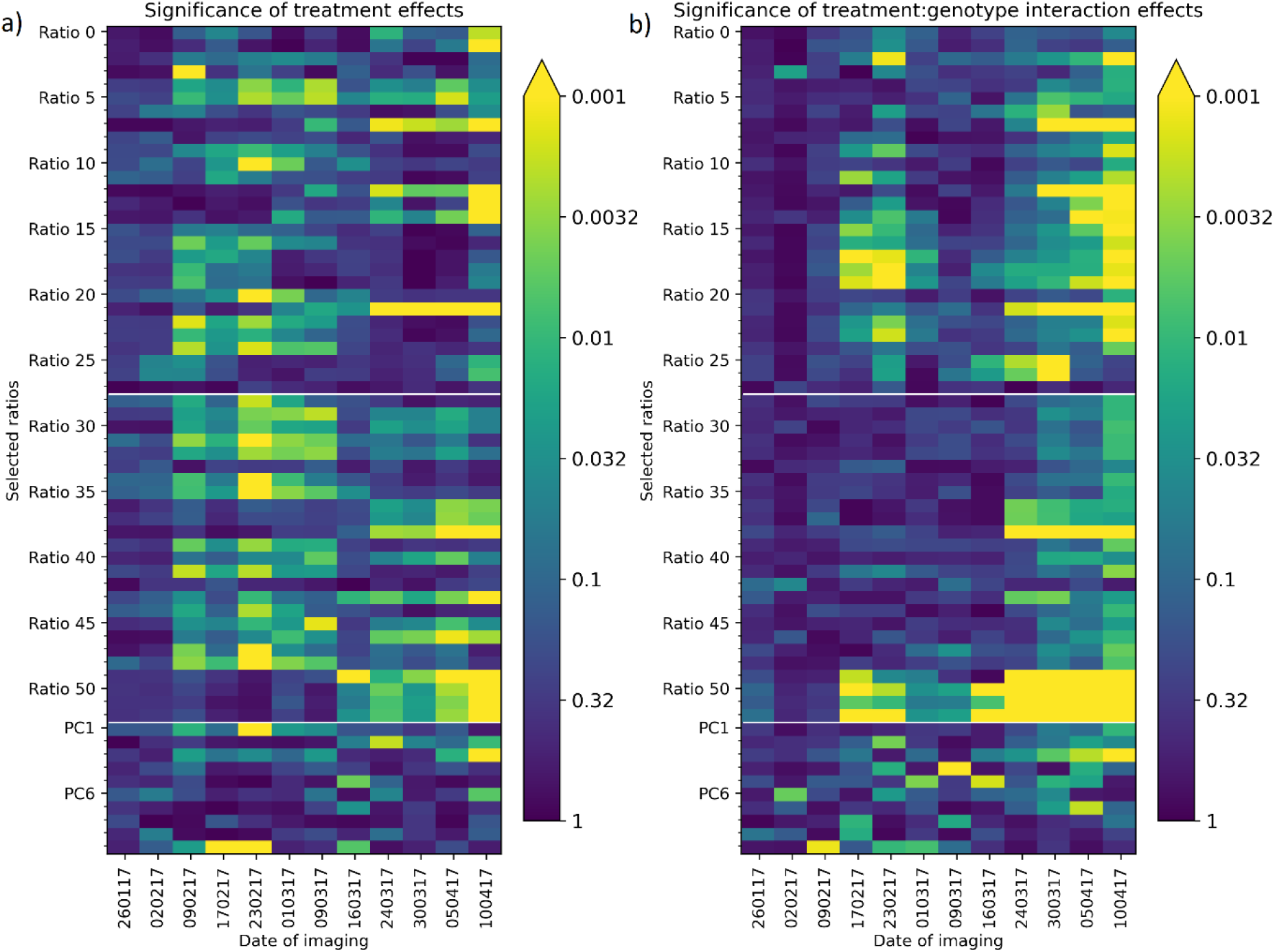
Heatmap plots showing (a) significant of treatment and (b) treatment*genotype interaction on the wavelength ratios and PC scores. p is plotted on a log scale and p<0.001 is used as the threshold to determine significance which is represented in the plot by the yellow colouring.

**Figure 6:**
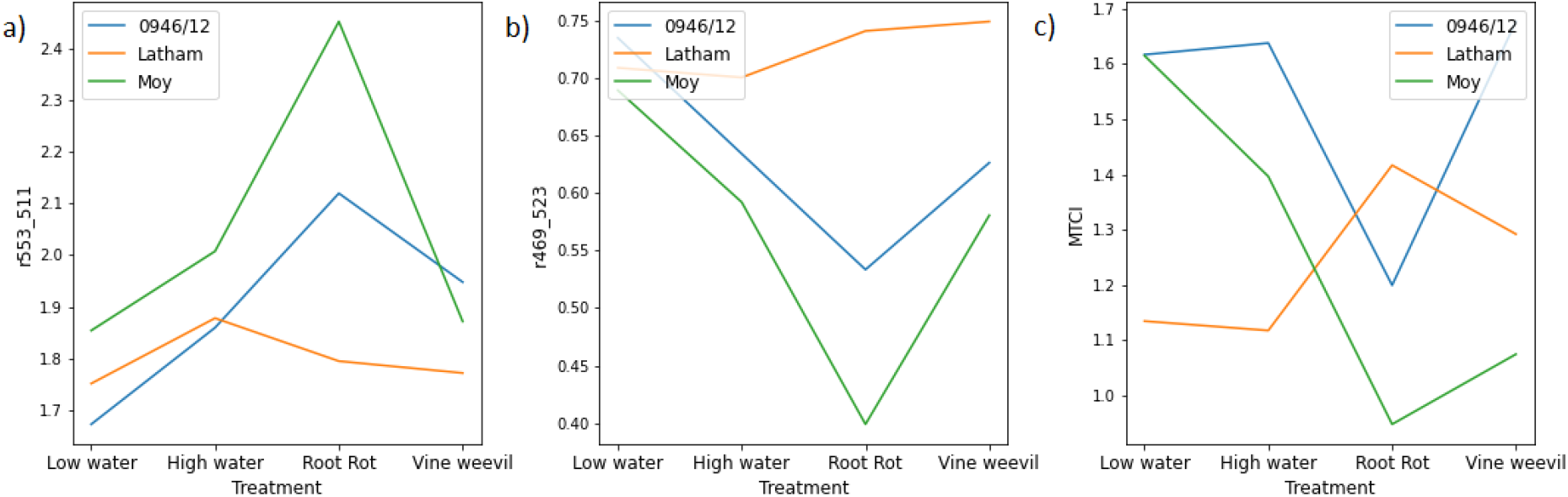
Plots of the response of individual wavelength ratios (a) r553_511, (b) r469_523 and (c) MTCI to the interaction between genotype and treatment the final day of imaging prior to plant harvest (10/04/17). Values are the mean of n=8 replicate plants. See Table 1 for more information about these wavelength ratios.

Three patterns were observed in the interactive effects of genotype and treatment on spectral data, shown in Figure 6 for data collected at the end of the experimental period (10/4/17). In some cases, the *P. rubi* treatment showed a strong positive or negative effect on spectral responses of Glen Moy and 0946/12 compared with Latham (Fig 6a, b). In other cases, Latham showed a positive spectral response to this treatment compared with the negative response of the other two genotypes, contrasting with the strong positive responses of the latter genotypes to the water and vine weevil treatments (Fig. 6c).

Supplementary figure 2 shows an example of temporal trends in the spectral response of each genotype to the four treatments for the ratio r469_523. In the *P. rubi* treatment, Glen Moy and, to a lesser extent, 0946/12 showed a reduction in response over time while Latham maintained a consistent response. A similar pattern was observed for the spectral response to the vine weevil treatment, but with smaller reductions in reflectance over time. For the two water treatments, the three genotypes showed comparable responses with little change in reflectance over time.

### Correlation between biophysical and spectral traits

We examined the correlation of wavelength ratios and PC scores with four biophysical measures: leaf number, leaf chlorophyll content, stomatal conductance, and soil moisture (figure 7).

**Figure 7:**
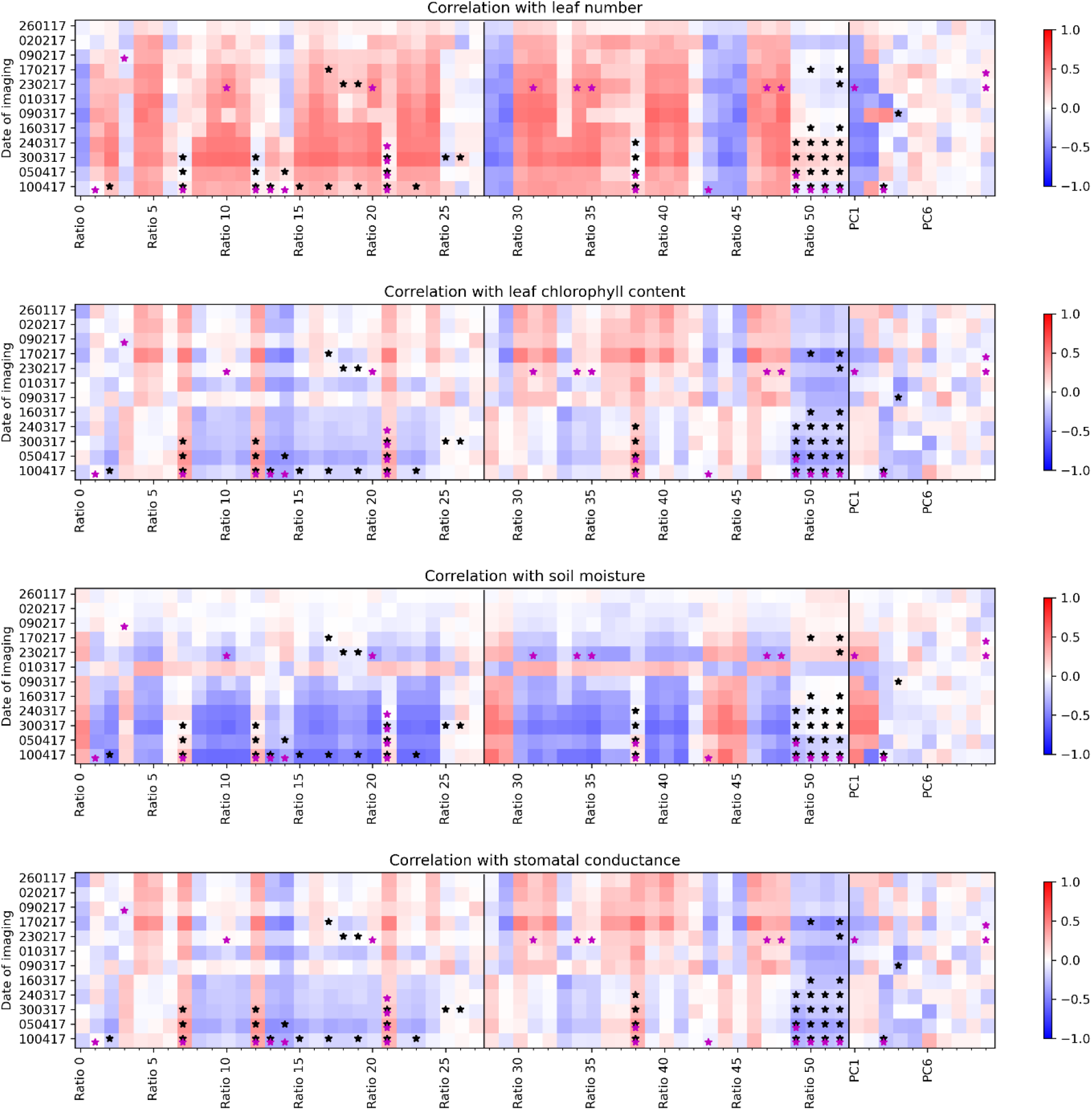
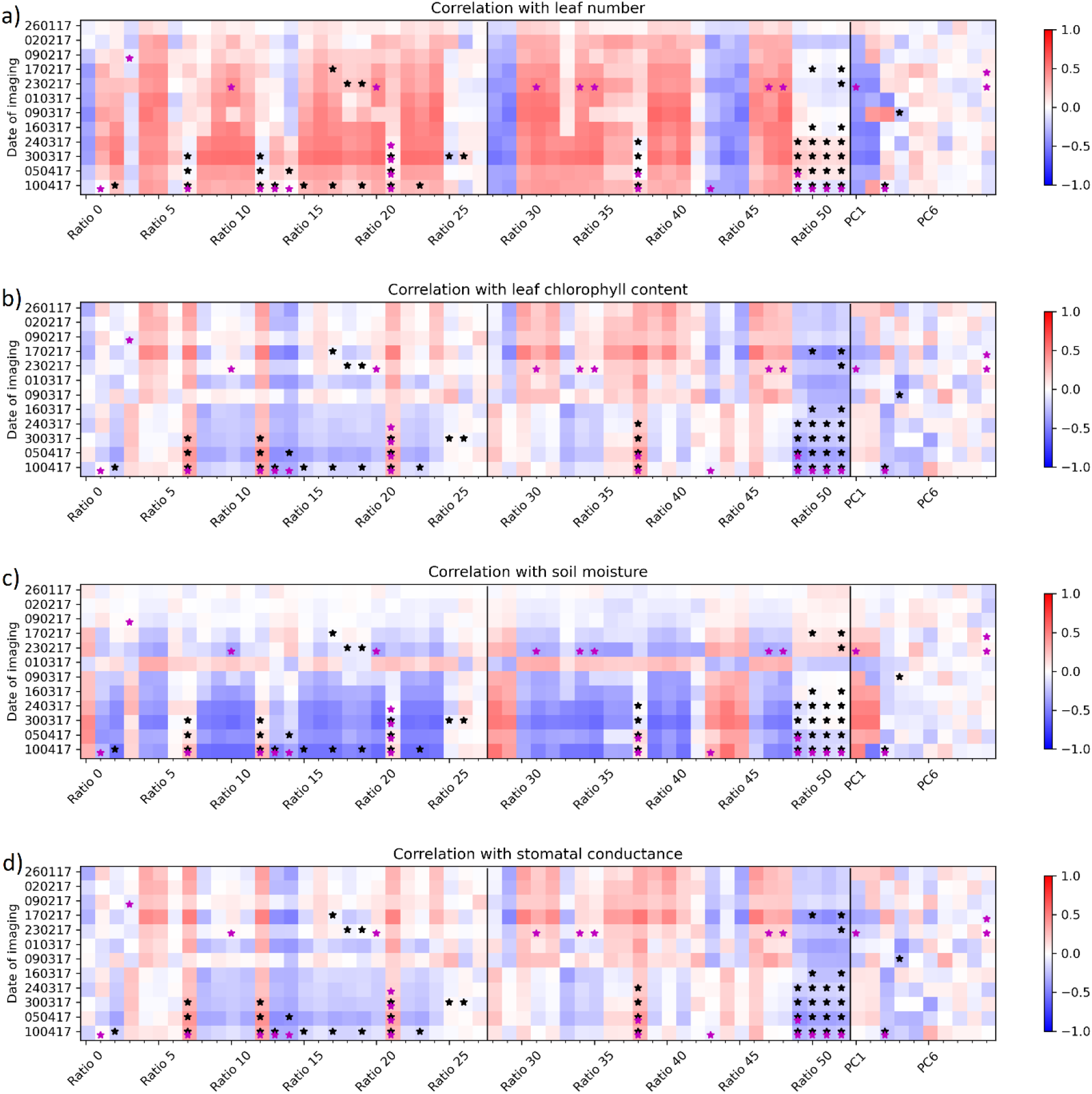
Heat maps showing positive or negative correlation of wavelength ratios derived from imaging data with (a) leaf number, (b) leaf chlorophyll content, (c) soil moisture, and (d) stomatal conductance data collected on the same date throughout the trial. Stars indicate significant differences for these ratios from reml analysis: black indicates significant treatment*genotype interaction and magenta indicates significant treatment effects. Genotype effects are not shown.

Several wavelength ratios correlated strongly with leaf number and chlorophyll content (fig. 7a, 7b). Ratio 10 or chl1green has strongest correlation with leaf number and ratio 39 or r719_691 has strongest correlation with chlorophyll content. There is no correlation between leaf number and chlorophyll content. These are similar ratios to the ones found to have significant interaction effects between treatment and genotype, indicating the treatment effect is due to changing levels of chlorophyll in the leaves and/or changes in the total reflective surface area (leaf number). By contrast, different wavelength ratios correlated with soil moisture (fig. 7c); these were weaker correlations than those detected for leaf chlorophyll content and leaf number, although the strength of the correlations increased during the experimental period. Correlations between spectral data and stomatal conductance were weak (fig. 7d). On 1/3/17 correlations for soil moisture are reversed, and the soil moisture data for this date was higher than normal and showed different treatment patterns compared with other dates. This is probably due to irrigation having been triggered just before soil moisture readings were taken resulting in unrepresentative measurements of soil moisture.

The correlations between wavelength ratios and harvest data are shown in figure 8. There were strong negative or positive correlations between most ratios and the vine weevil damage score and root mass. Moderately strong correlations were detected between many wavelength ratios and above ground growth metrics, particularly leaf mass. Root length and vine weevil larvae count did not correlate with any of the tested ratios. This shows that many of the wavelength ratios can indicate the relative response of above- and below-ground plant traits to stress treatments that can only be measured by destructive harvesting. Scatterplots showing the relation between selected wavelength ratios and plant biophysical data (figure 9) indicated linear (fig. 9a, c) and nonlinear (fig. 9b,d) relations between the variables.

**Figure 8:**
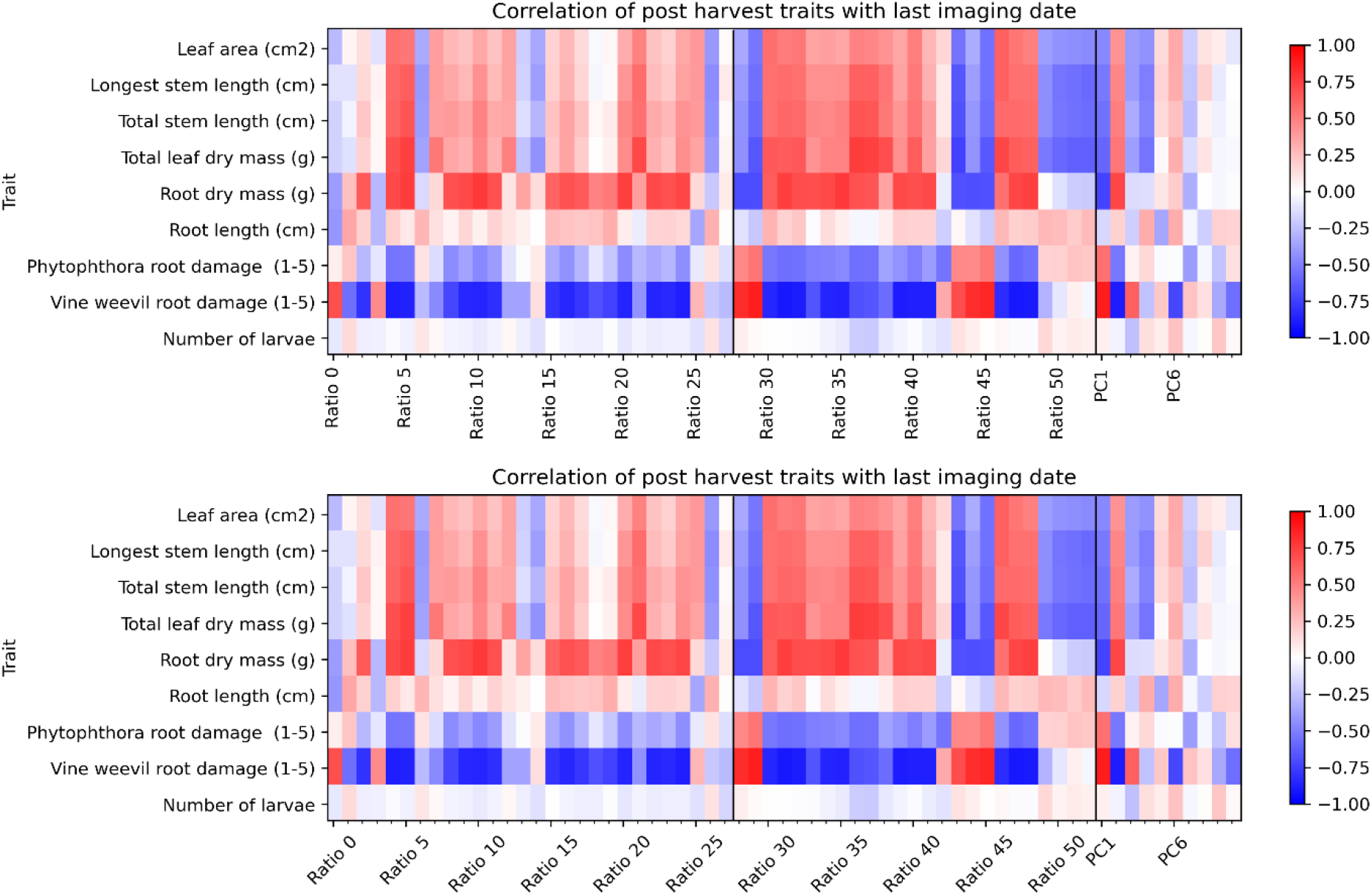
Heat map showing positive or negative correlation of wavelength ratios and principal components derived from imaging data collected 2 days before harvest with biophysical traits measured at plant harvest.

**Figure 9:**
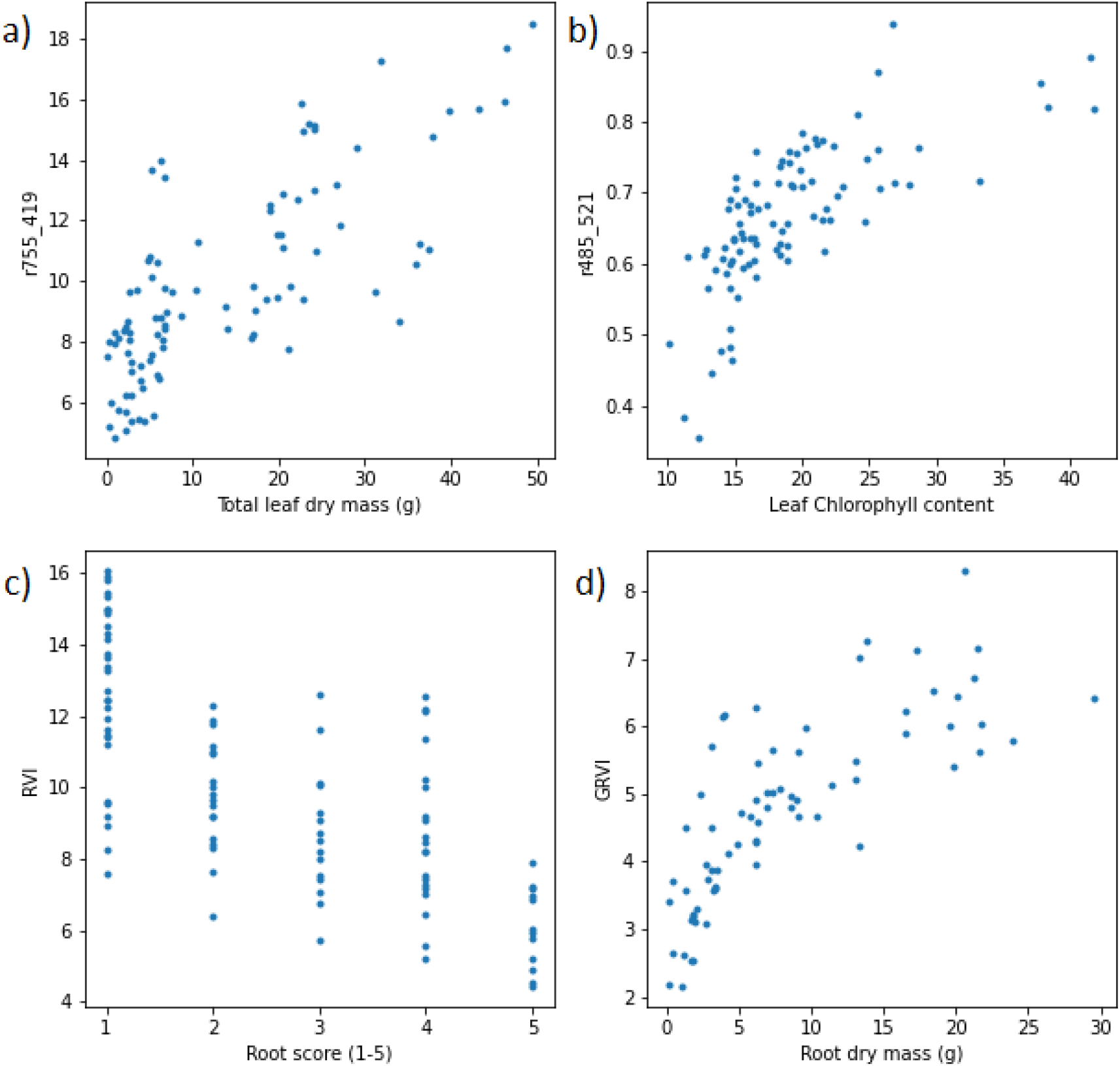
Scatterplots showing the relation between intensity of selected wavelength ratios and (a) total leaf dry mass, (b) leaf chlorophyll content, (c) root damage score and (d) root dry mass quantified prior to or at plant harvest.

## Discussion

The development of methods for non-destructive assessment of plant stress responses is crucial for overcoming bottlenecks associated with high throughput phenotyping of genotypes for crop improvement and rapid monitoring of plant stress in crop plantations. Here, we have demonstrated that hyperspectral imaging can detect differences between abiotic and biotic stress treatments in raspberry and identified wavelength ratios showing differential genotypic responses to these stresses. Some of these spectral signatures correlated strongly with plant growth and biophysical traits indicating they could act as non-destructive indicators of plant stress. Table 2 below summarises the most important spectral measures detected in this study that correlate with the different biophysical traits or show specific treatment responses.

**Table 2:** Table showing selected ratios picked out in this study as best for particular tasks. Ratios 36, 39 and 49-52 are novel to this work and outperformed existing ratios.

| Thing measuring | Useful ratio numbers | Ratios | Definitions |
| --- | --- | --- | --- |
| Distinguishing treatment | 21 | SR4 | R700/R670 |
| Treatment*genotype interaction | 49 | r509_512 | R509/R512 |
|  | 50 | r568_641 | R568/R641 |
|  | 51 | r557_658 | R557/R658 |
|  | 52 | r537_644 | R537/R644 |
| Leaf Number correlation | 10 | ch1green | R800/R550 – 1.0 |
| Soil Moisture correlation | 39 | r719_691 | R719/R691 |
| Chlorophyll content correlation | 51 | r557_658 | R557/R658 |
| Root dry mass correlation | 35 | GRVI | nir/green |
| Leaf dry mass correlation | 36 | R753_417 | R753/R417 |

### Raspberry genotypic responses to stress

The three genotypes were selected based on known differences in plant characteristics, primarily based on their responses to *P. rubi* infection, but also differences in plant stature and canopy density. Glen Moy was chosen as it is a susceptible variety while Latham and 0946/12 have both previously shown resistance to *P. rubi* infection. Resistance to root rot in Latham is thought to relate to more vigorous root growth than in susceptible Glen Moy^[39]^ and this might also contribute to varietal differences in vine weevil susceptibility^[40]^. In this study, Latham and Glen Moy both had the lowest root damage score in the *P. rubi* treatment indicating that the infection was not the main factor affecting root growth. A similar pattern was seen in above ground measures with canopy height being tallest in the *P. rubi* treatment and shortest in the low water treatment.

Canopy height of the three genotypes responded differentially to the imposed treatments, which was driven by differing responses to *P. rubi* treatment, for which 0946/12 and Glen Moy plants were the tallest. By contrast, Latham plants were taller in the high water and vine weevil treatments.

In the spectral data, ratios with the largest genotype* treatment interaction effect showed the biggest genotypic differences in the *P. rubi* treatment. Latham showed a different response to the other two genotypes in several ratios that have been previously linked to leaf chlorophyll content (e.g. GI r554_R667^[24]^). This supports the conclusion that we are picking up a measure of plant response to the *P. rubi* infection on the plants.

### Using spectral data to differentiate genotypic responses to stress

Using spectral data to detect differential responses of raspberry genotypes to the stress treatments identified three patterns. Of the wavelength ratios showing significant interactions between treatment and genotype, two groups of spectral traits showed genotypic differences in their response to the *P. rubi* treatment (positive or negative change in ratio intensity) while a third group of spectral traits also showed differential responses to the vine weevil treatment. These patterns indicate that the ratios can detect different plant physiological responses to stress. Previous studies using similar ratios have linked variation in reflectance intensity to changes in chlorophyll content^[24,26]^, which in the present study might relate to either changes in leaf chlorophyll content or plant leaf density. Newer leaves will generally have a lower chlorophyll content compared to older ones so a reduction in leaf chlorophyll content could be indication of a more rapidly growing plant.

In the *P. rubi* treatment, ratios showing large responses mostly included wavelengths in the green area of the spectrum, while those responding to the vine weevil treatment included the red edge area of spectrum. Some of these ratios have been previously shown to detect reflectance changes at high chlorophyll concentrations where other ratios were saturated^[19,32]^. Both leaf chlorophyll concentration and leaf mass responded significantly to the interactive effects of genotype and treatment at the end of the experiment: in the *P. rubi* treatment, leaf chlorophyll content was reduced for 0946/12 and Glen Moy but increased in Latham, while leaf mass showed the reverse trend. In addition, in the vine weevil treatment leaf mass was reduced for Glen Moy compared with the high-water treatment but was similar between the two treatments for Latham and 0946/12. The finding that plant spectral traits co-varied with several plant biophysical traits in response to belowground biotic stress suggests that spectral traits could act as indicators of plant stress.

What is not clear, however, are the underlying mechanisms leading to changes in leaf reflectance characteristics in response to belowground biotic stress. Previous work has shown typical symptoms of *P. rubi* are a reduced number of lateral shoots, leaves turning yellow, plants wilting and generally stunted growth ^[43]^. Although classic root rot symptoms were not observed in this study, there were clear effects of *P. rubi* on canopy height and leaf mass, which were detected in spectral data. Previous work looking at vine weevil infestation found that increased vine weevil presence caused reduced plant growth with significant reductions in plant mass^[40]^. Positive correlations between number of vine weevil eggs and leaf nitrogen content have been previously reported which could relate to increased chlorophyll concentration in infested plants due to reduced production of new leaves^[44]^. It is worth noting that precise measurement of biophysical plant responses to below ground stress is difficult, and the biophysical measurements reported here are also imperfect proxies for stress. Further work observing in more detail the plant responses to insect attack, pathogen infection or water deprivation and an attempt at calibrating hyperspectral data to plant physiological parameters would be helpful in refining the use of hyperspectral imaging for phenotyping plant stress responses.

#### Further development of hyperspectral imaging as a stress detection tool

This study demonstrates that hyperspectral imaging can be used to detect plant responses to stress and differential responses of crop genotypes. This opens the potential of imaging to be used for plant stress detection. There was, however, limited evidence that our current method can distinguish different causes of plant stress, whether due to root pests or pathogens or due to water limitation. This may be due to our image segmentation method which currently treats each plant as a single sample point. A finer grained segmentation method that was able to distinguish different leaves on the same plant may be able to generate more information about the spatial distribution of symptoms in a plant which would provide further information on plant stress status.

Spectral responses to plant stress varied during plant growth and varied between genotypes, which has implications for how spectral data should be collected and used for different purposes. It should be noted, however, that temporal changes in spectral reflectance may be due in part to imperfect image normalisation. The lighting environment in the glasshouse was not entirely controlled, and although high-pressure sodium grow lamps were turned off during imaging, natural light was supplemented with halogen lamps to illuminate the plants when imaging was carried out. We attempted to control for variations in incident light using a white reference standard tile. The spectrum of each image was normalised using reflectance from this normalisation tile which in theory converts absolute intensity to relative reflectance values. However, the 3-dimensional structure of the plants and directional nature of both natural sunlight and additional halogen lamps means that this was not a perfect process. Differences in reflectance from day to day could be partly due to different weather conditions or different sun position in the sky which would change the observed reflectance spectrum. Solutions to this issue could involve carrying out imaging under more controlled lighting conditions or using an improved reflectance reference which has 3D structure.

### Conclusions

Overall, we have shown that hyperspectral imaging can be used to distinguish differences in both raspberry plant responses to stress treatments and differential stress responses between genotypes. We have found spectral reflectance ratios that we can link to biotic below ground stresses experienced by the plant. Further work is required to understand the physiological processes underpinning spectral changes in these ratios and how they respond to the imposed stress. This study has shown that spectral imaging can be used for plant phenotyping and stress monitoring as it is able to detect differences between genotypes in their responses to imposed plant stress.

## Acknowledgements

The authors would like to thank Dr Ankush Prashar (University of Newcastle) and Prof Lyn Jones (University of Dundee) for their advice on set up of imaging platform and experimental design. We would also like to thank former college Dr Christine Hackett (retired/Biomathematics & Statistics Scotland) for statistical advice and support in analysing the data. This work was supported by Innovate UK (grant No. 101819), the strategic research programme (2016-2022) and the Underpinning Capacity project ‘Maintenance of Insect Pest Collections, both funded by the Rural and Environmental Science and Analytical Services Division of the Scottish Government.

## Author Contributions

DW and AB collected the data. DW processed the image data. DW, AK and JG interpreted the results. DW drafted the manuscript. JG and AK significantly edited the manuscript. All authors read and approved the final manuscript.

## Data Availability

The data supporting the findings of this study are available from the corresponding author, Dominic Williams, upon request.

